# Identification and analysis of DEGs of mouse tumor cells by integrated bioinformatics methods

**DOI:** 10.1101/2024.12.24.629056

**Authors:** Xizhong Jing, Baiyong Li, Yifei Wang

## Abstract

Mutations and genome instability are hallmarks of cancer. The advanced Next Generation Sequencing(NGS) technology now enables rapid and cost-effective analysis for tumoral and normal genomic translational research. In this research, mRNA sequencing was applied to get the whole transcriptome data of mouse B16F10 melanoma and CT26 colon carcinoma cells. Up regulation genes and down regulation genes were identified in tumor cells with differentially expressed genes(DEGs) analysis. The DEGs were analyzed with GO, KEGG and Reactome pathway enrichment study. Results demonstrated the DEGs and enriched pathways were mainly associated with cell proliferation and biosynthetic processing. Further, gene fusion and alternative splicing analysis reveled fusion genes and skipped exons were extensive exist within the chromosomes of tumor cells. Together, the integrated DEGs and bioinformatics analysis in this article may shed light on further research of cancer treating targets and molecular mechanisms of carcinogenesis.

## Introduction

The genome and transcriptomic instability are hallmarks and prominent characteristics of cancer^[1]^. The genetic mutations that contribute to the tumorigenesis have been the subject of extensive research. Some driver mutations have the potential to transform a normally functioning cell into a cancer one^[2]^. The nucleotide sequence of DNA and some chromosomal abnormalities are heritable changes of cancer cells. Likewise, the transcriptomic profile of cancer is the set of RNA molecules expressed by tumor cells under certain conditions^[3]^. It varies based on the biological process, stage of cancer development, and microenvironment of the tumor^[4]^. NGS technology coupled with bioinformatic approaches has been widely used to reveal the processing of tumor cellular transcription and metastasis^[5]^. Now with the development of rapid and relatively inexpensive technologies for DNA and mRNA sequencing, including whole exome sequencing(WES), whole genome sequencing(WGS) and genome-wide association studies(GWAS)^[6,7,8]^, the identification of the DEGs has been greatly facilitated. These sequencing and analysis technologies now enable to identify the subtle mutations for prognostic and diagnostic markers as well as therapeutic targets^[9,10,11]^. Analyzing the transcriptome of tumor and normal cells can help us for a better understanding of the molecular mechanisms of carcinogenesis. Most cancer research starts out with the investigation of the biological and genetic differences between normal cells and cancerous cells. High-throughput transcriptome mRNA sequencing (RNA-Seq) has become the main option for DEGs analysis^[12,13]^. RNA-seq technology has some advantages over the DNA sequencing, such as the high level of data reproducibility and allowing to identify and quantify the expression of isoforms and unknown transcripts^[14,15,16]^. By studying the DEGs between tumor cells and normal tissue cells, researches can discover the mechanisms by which cancer cells grow, transform from normal cells into cancer cells, and metastasize throughout the body^[17]^.

Cancer cell lines are the main workhorse of cancer research^[18]^. Generally, cancer cell lines are clonal and genetically stable thus the results obtained from one laboratory can be readily extended to another. It is known that cell lines are to evolve in culture and become resultant genetic and transcriptional heterogeneity^[19]^. Studies of the evolution of different cell lines can lead to insights into cancer progression and initiate the development of new clinical treatments^[20]^. Further, investigations into the different cell signaling pathways can reveal molecular alterations that are shared among different types of cancers and develop possible strategies for therapy. B16F10 melanoma cells and CT26 colon carcinoma cells are widely used for establishing preclinical tumor models and exploring potential cancer therapy. B16F10 cells are characterised with weak intrinsic immunogenicity and high metastatic potential^[21]^. Murine melanoma model can be set up by inoculating B16F10 cells in syngeneic C57BL/6 mouse strain. The intrinsic properties of B16F10 cells induce a moderate host antitumor immune response in C57BL/6 mice. CT26 cell line is a colon carcinoma cell line derived from BALB/c mouse. CT26 tumor models share similar molecular features with aggressive, undifferentiated, refractory of human colorectal carcinoma^[22]^. In this study, we extracted the mRNA from B16F10 and CT26 tumor cells as well as normal mouse tissues and applied RNA-Seq for DEGs analysis. We aim to find novel tumor targets and molecular mechanisms of carcinogenesis.

## Results

### RNA-seq & Sequencing alignment data quality control

To get the gene expression of tumor cells and normal mouse tissues, the whole transcriptomic sequencing was applied. Total mRNA was extracted from B16F10 and CT26 tumor cells as well as normal mouse tissues. Double-stranded cDNA was synthesis for sequencing library establishment. The illumina HiSeq2000 system was used for gene expression analysis. The detailed workflow of of RNA-seq experiments was shown in figure 1a. The bioinformatics analysis process of mRNA transcriptome sequencing mainly includes data quality control, raw data alignment with reference genome, gene expression analysis, DEGs analysis, functional enrichment analysis, fusion genes and alternative splicing analysis. The workflow for identification and analysis of DEGs was shown in figure 1b.

**Figure 1.**
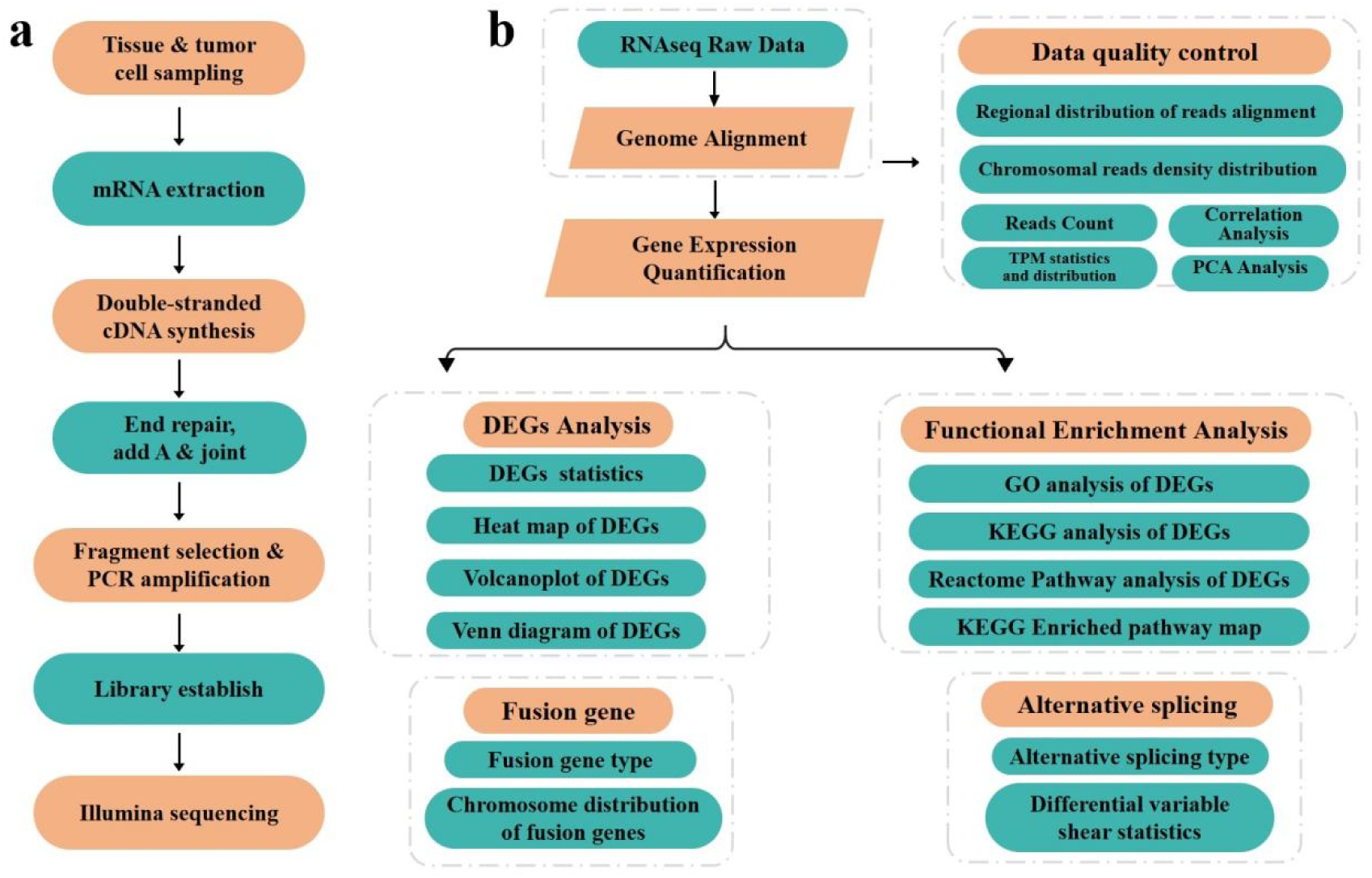
The workflow of RNA-seq & integrated bioinformatics analysis processing. a, Workflow of of RNA-seq experiments. b, Bioinformatics analysis processing.

HTSeq was used to analysis the sequencing aligned gene reads. The distribution of aligned reads on chromosomes was shown in figure S1a. Most sequencing reads were distributed on the exon and CDS regions as shown in figure S1b. Only a small fraction of aligned reads were distributed on 5’UTR and 3’UTR. These results indicated the RNA-seq aligned gene reads were mainly responsible for gene coding. TPM (Transcripts Per Kilobase of exon model per Million mapped reads) divides the read counts by the length of each gene in kilobases^[23]^. The TPM distribution in figure S1c demonstrated the consistent of gene expression of tumor cell lines and normal tissues under experimental conditions. Correlation analysis is a statistical method to evaluate the relationship between experimental samples. The similarity and difference between different experimental conditions can be observed through correlation analysis for non-biological replicates samples. In figure S1d, the different samples correlation was shown in different color. In figure S1e, Principal Component Analysis(PCA) was applied to compare the similarities and outliers of the tumor cells and normal tissue samples. Results indicated the comparable relationship of the B16F10 and CT26 tumor cells and normal mouse tissue sequencing samples.

### Functional enrichment analysis of DEGs of tumor cells

We next investigated the reads number of DEGs in tumor cells and normal tissue samples. In figure2 a-d, the red color represents the up-regulation genes and green represents down-regulation genes. More up-regulation genes were enriched in tumor cells compared with normal tissues in heat map and volcanoplot. We identified 5870 up regulation genes and 2292 down regulation genes in B16F10 cells. For CT26 cells, 7714 upregulated genes and 2665 downregulated genes were identified respectively. We next used Wayne study to analysis the DEGs distribution. 16370 genes were co-expression by normal C57BL/6 tissues and B16F10 cells. 8778 and 2319 genes were expressed by normal C57BL/6 tissues and B16F10 cells respectively. For BALB/c and CT26 cells samples, the co-expression gene number were 16999. 6177 and 5090 genes were only expressed by BALB/c tissue cells and CT26 cells, respectively (figure 2 e-f). The detailed DEGs information is listed in supplementary materials.

**Figure 2.**
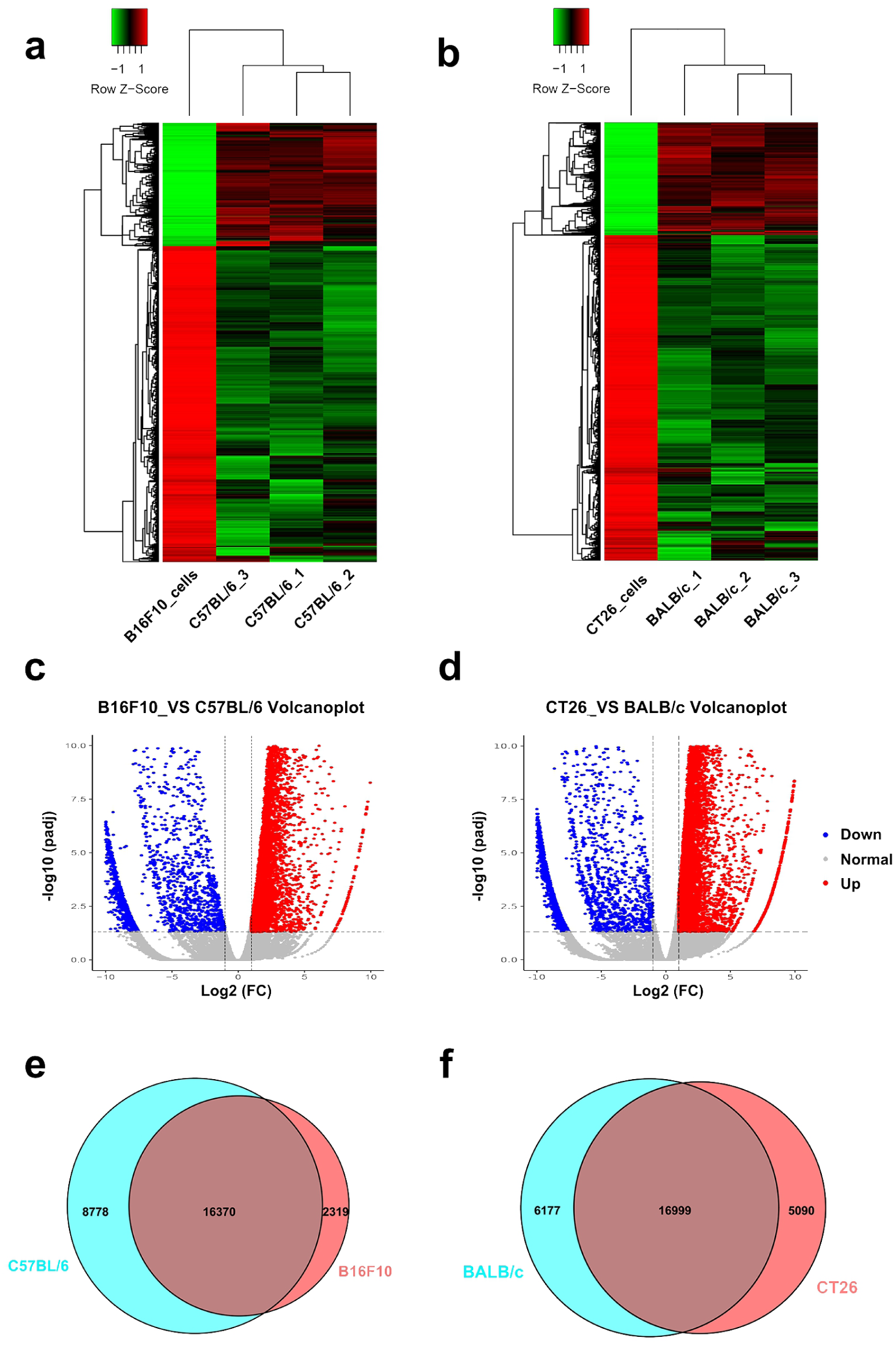
Differentially expressed genes (DEGs) analysis. a, Heat map of B16F10 cells and C57BL/6 mice DEGs clusters. b, Heat map of CT26 cells and BALB/c mice DEGs clusters. c, Volcanoplot of B16F10 and C57BL/6 mice DEGs. d, Volcanoplot of CT26 cells and BALB/c mice DEGs. e, Venn diagram of B16F10 and C57BL/6 mice DEGs. f, Venn diagram of CT26 cells and BALB/c mice DEGs.

To obtain a further insight into the biological functions of DEGs, GO annotation, KEGG and Reactome pathway enrichment analyses were performed (figure 3–5). The enriched top10 GO terms were shown in figure 3 a & b. The GO terms were comprised of 3 parts: biological process (BP), cellular component (CC), and molecular function (MF). DEGs of BP were involved in ameboidal type cell migration, autophagy, chromosome segregation, mRNA processing, mitotic cell cycle phase transition and regulation of cellular macromolecule biosynthetic process. DEGs of CC were involved in actin cytoskeleton, asymmetric synapse, cell leading edge, chromosomal region, collagen-containing extracelular matrix, microtubule, nuclear envelope, nuclear speck, postsynaptic specrialization and spindle. DEGs of MF were involved in GTPase regulator activity actin binding, RNA catatytic activity, Nucleoside-triphosphatase regulator activity, phosphoipid binding, protein serine/threonine kinase activity, Transcription coregulator activiy and ubiquitin-like protein transferase activity. Besides, the enriched KEGG pathways as presented in figure 4, including cell cycle, focal adhesion, proteoglycans in cancer, Salmonella infection, Human papillomavirus infection, Regulation of actin cytoskeleton, Human T-cell leukemia virus 1 infection, PI3K-Akt signaling pathway, Amyotrophic lateral sclerosis, Pathways of neurodegeneration-multiple diseases, Parkinson disease, MAPK signaling pathway, Alzheimer disease, Ras signaling pathway, Human cytomegalovirus infection and Prion disease pathway.

**Figure 3.**
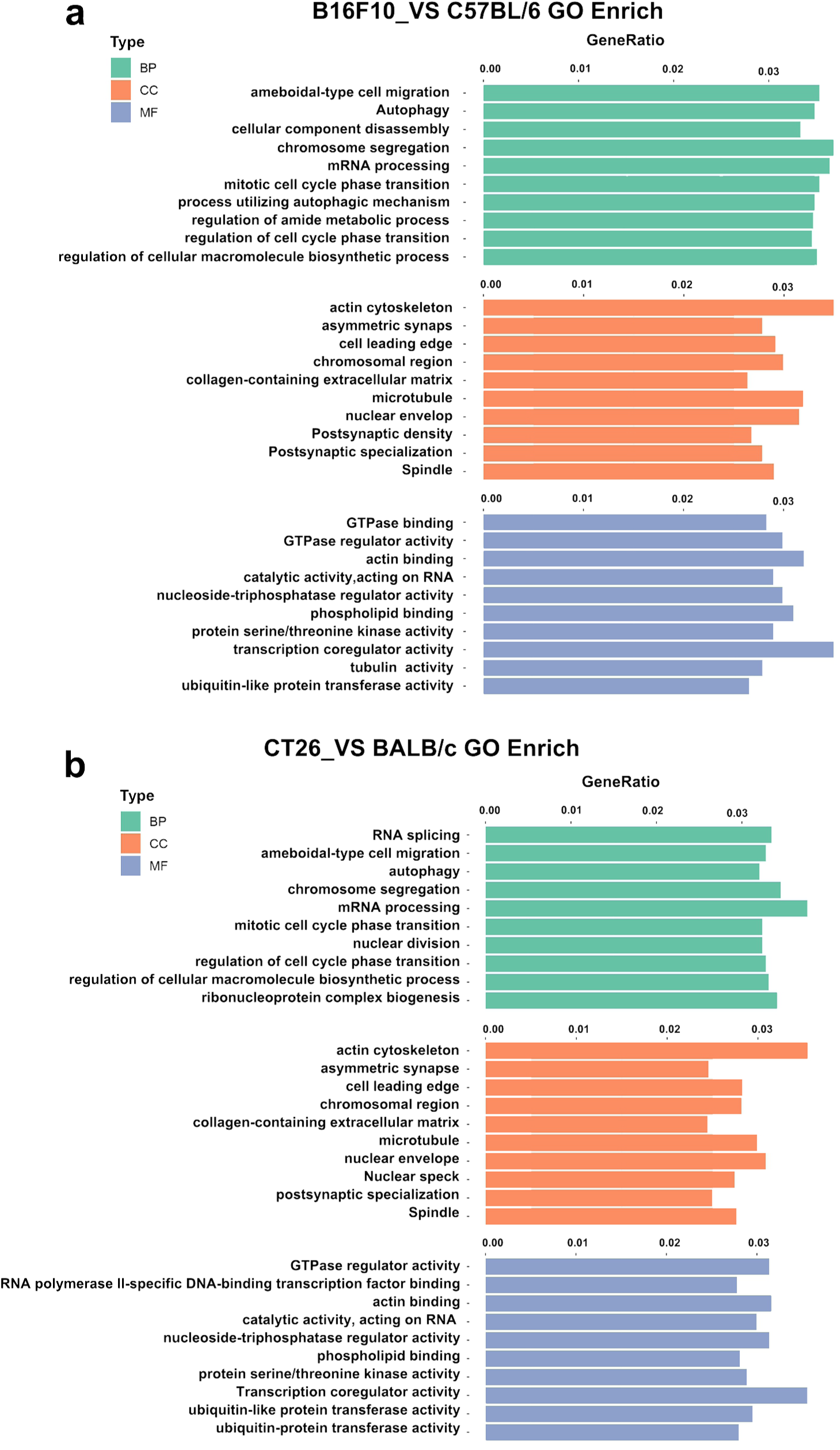
GO enrichment analysis of DEGs. DEGs: differentially expressed genes; GO: Gene Ontology. (P-value<0.01 and q-value<0.05)

**Figure 4.**
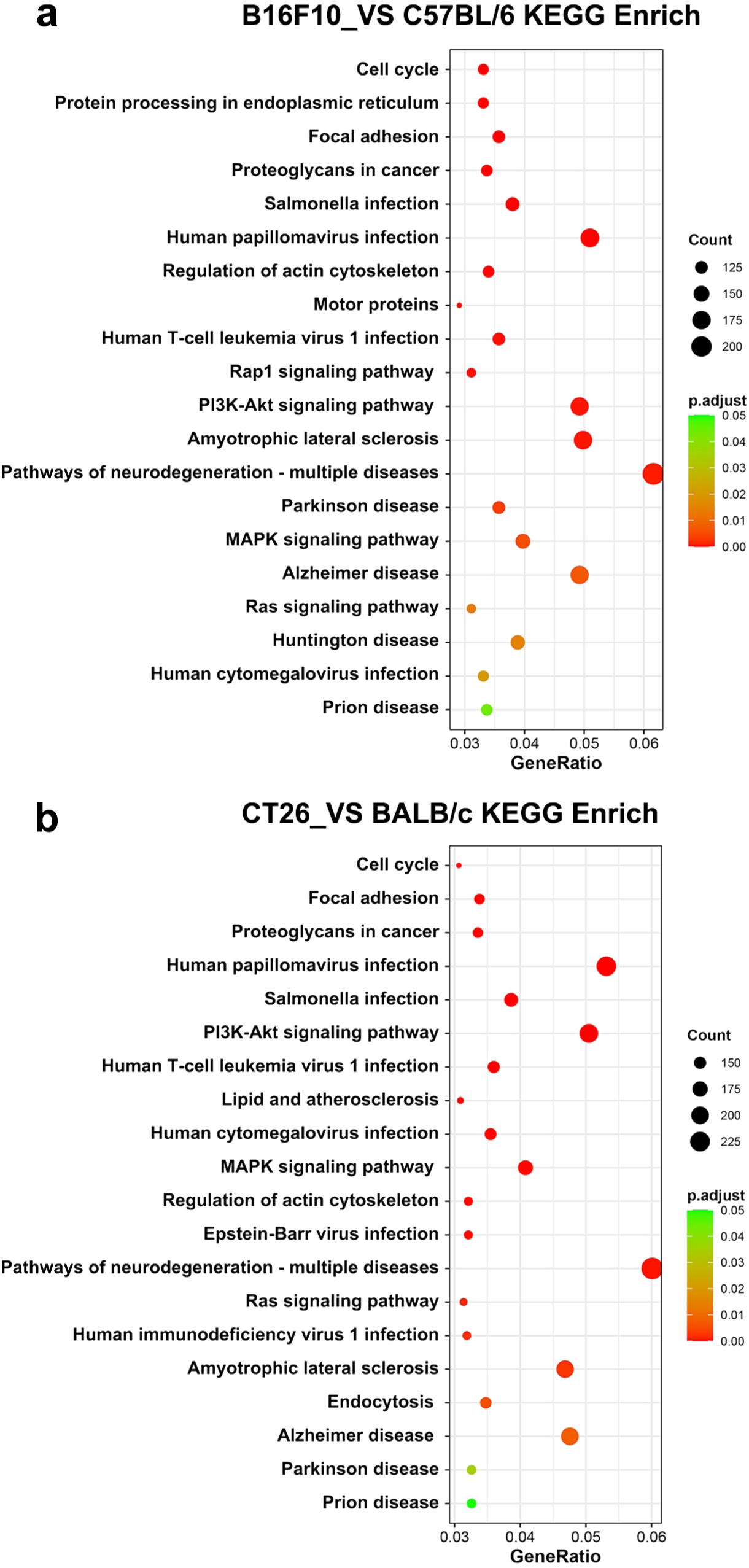
KEGG enrichment analysis of DEGs. DEGs: differentially expressed genes; KEGG: Kyoto Encyclopedia of Genes and Genomes.(P-value<0.05 and q-value<0.05)

**Figure 5.**
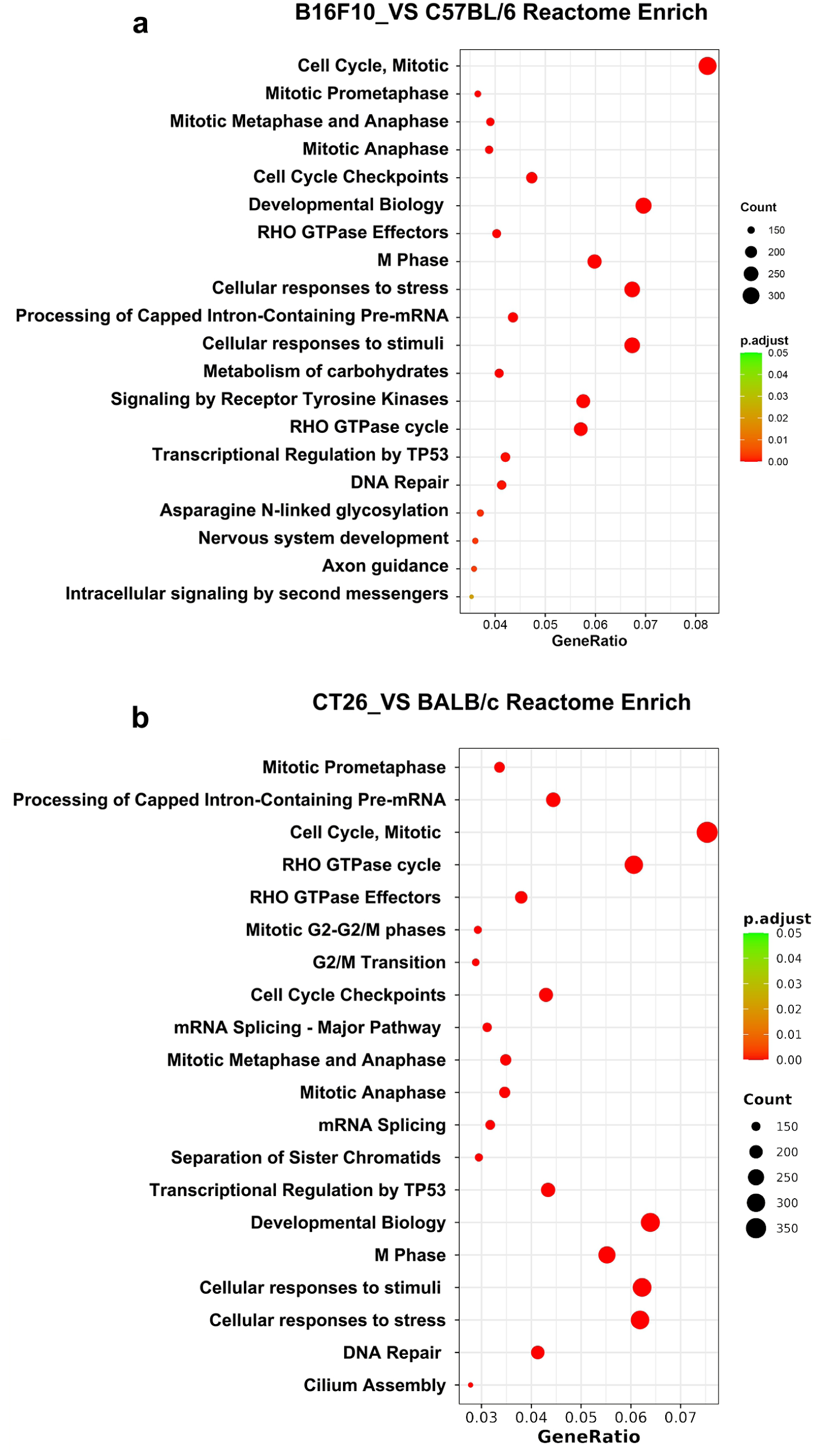
Reactome pathway enrichment analysis of DEGs. DEGs: differentially expressed genes.(P-value<0.05 and q-value<0.05)

Further, Reactome database^[24]^ (http://www.reactome.org) mainly annotates cellular component, including proteins, nucleic acids, lipids, carbohydrates for their biology processes and records the function of proteins associated with disease. The clusterProfiler hypergeometric distribution algorithm^[25]^ was applied to investigated the enrichment of Reactome pathways of DEGs. The manly reactome pathways associated with B16F10 and CT26 tumor cells were cell cycle and DNA repair, including cell cycle mitotic, cell cycle checkpoints, RHO GTPase effects, cellular response to stress & stimuli, developmental biology, DNA repair and axon guidance (figure 5). Taken together, these GO terms, KEGG and Reactome pathways analysis demonstrated the DEGs of B16F10 cells and CT26 cells were associated with cell cycle and cellular replication biological processes in normal function and diseases.

Since cell cycle pathway is highly associated with the enrichment analysis of DEGs, we also investigated the expression genes in cell cycle pathway. The significantly enriched DEGs of B16F10 and CT26 cells were shown in figure 6–7. The up-regulated and down-regulated genes were marked with red and green color, respectively. It is no wonder that more up-regulated genes with cell cycle were enriched in B16F10 and CT16 cells than down-regulated genes. Very small number of down-regulated genes were enriched and the expression pattern of the down-regulated genes in the two cell lines are quite similar. For example, Cdkn2b and Cdkn1c are negative checkpoint of cell proliferation in G1 phase of cell division^[26]^. Our results showed the two genes were down-regulated in both B16F10 and CT26 cells (figure 6–7). Similar, Sulforaphane(Sfn) is a S-phase checkpoint of cell cycle^[27]^ and down-regulated in both two cell lines (figure 6–7). However, Gadd45a gene was down-regulated in B16F10 and up-regulated in CT26 cells. TGFβ2 gene was up-regulated in B16F10 and down-regulated in CT26 cells (figure 6–7). These results indicated slight differences exist in the cell cycle expression patterns of the two tumor cell lines. Other enriched pathway maps are included in supplementary materials.

**Figure 6.**
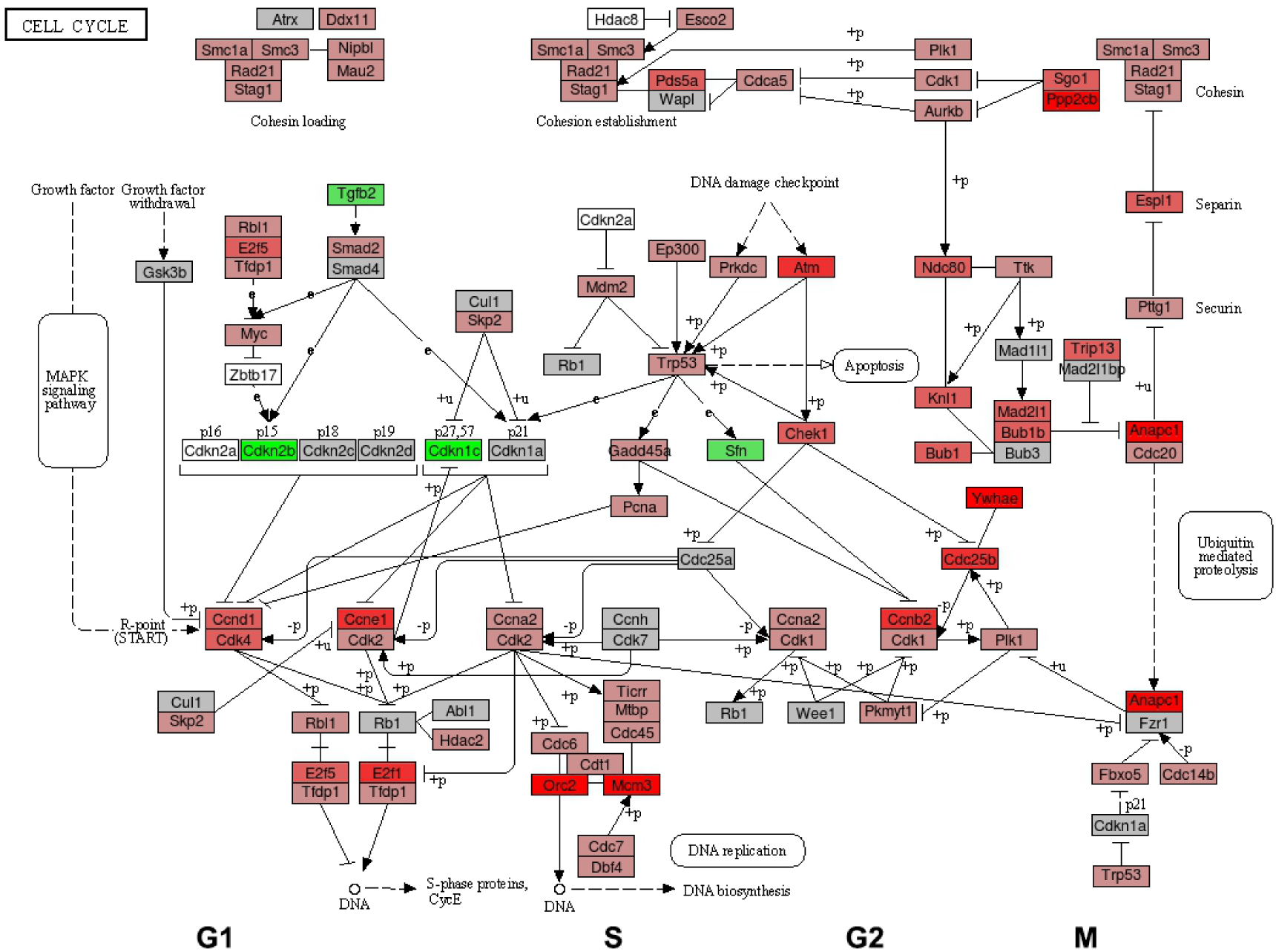
KEGG enriched DEGs of B16F10 cell cycle pathway map. DEGs: differentially expressed genes; KEGG: Kyoto Encyclopedia of Genes and Genomes.

**Figure 7.**
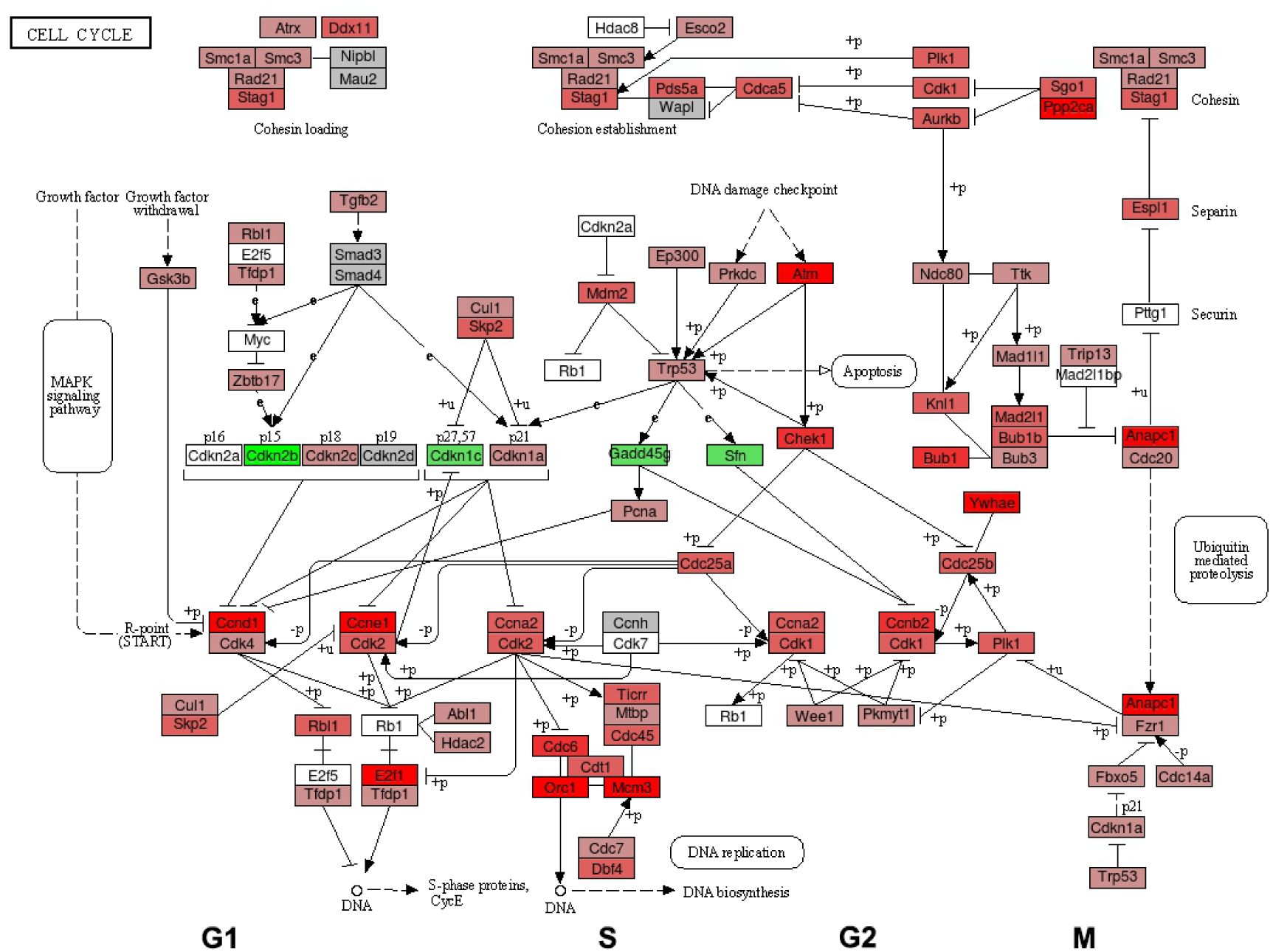
KEGG enriched DEGs of CT26 cell cycle pathway map. DEGs: differentially expressed genes; KEGG: Kyoto Encyclopedia of Genes and Genomes.

### Fusion genes and alternative splicing analysis

Gene fusion is fusion of part or all of the gene sequences (promoters, enhancers, ribosomal binding sequences, terminators, *etc*.) in separated genes^[28]^. Gene fusion can occur at the transcriptome level. mRNA transcribed from two different genes for some reason fused together to form a new fusion mRNA, which may encode proteins or not. Fusion gene is a hallmark of many types of tumors^[29]^. To have a more better knowledge of the location of genes fusions in chromosomes, we used START-FUSION software to investigated the the distribution of fusion genes in the whole chromosomes (figure S2). We identified more fusion genes were existed within the chromosomes of B16F10 and CT26 tumor cells than normal tissue cells. In figure 8, the blue curve indicated the exist of gene fusion within one chromosome, and the orange curve indicated gene fusion in two chromosomes. For B16F10 cells, fusion genes within one chromosome mainly exist in chromosome 4, 5, 7, 8, 10, 12 and 17. Fusion genes in two chromosomes mainly existed in chromosome 2&7, 2&9, 2&15, 4&10, 6&13, 9&19, 11&17. For CT26 cells, fusion genes within one chromosome mainly existed in chromosome 2, 4, 5, 6, 8, 13, 16, 18 and X. Fusion genes in two chromosomes existed in chromosome 2&5, 2&9, 5&15, 14&17 (figure 8 a & b). The detailed fusion genes of B16F10 cells and CT26 cells were listed in Table 1 & Table 2.

**Figure 8.**
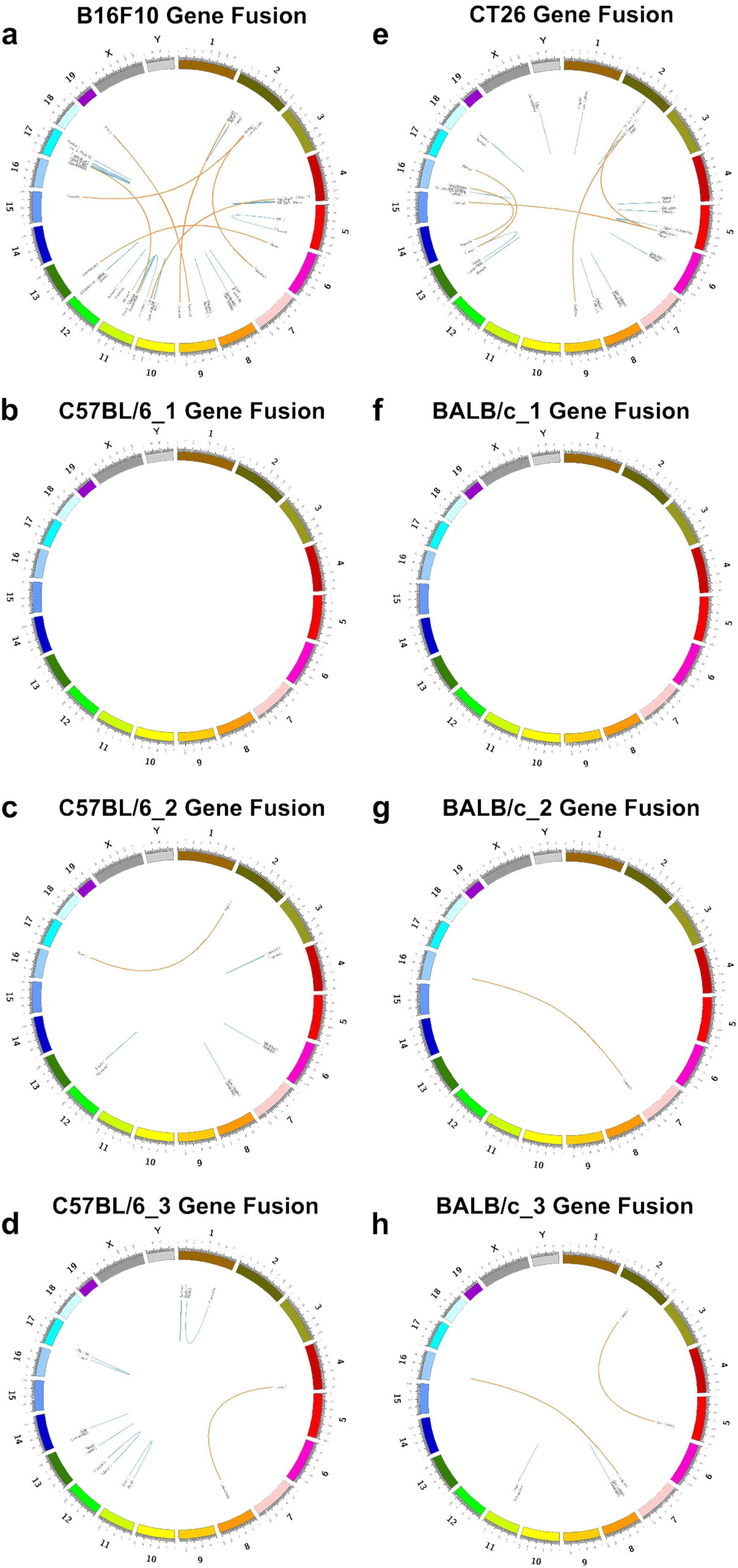
Distribution of fusion genes in chromosomes. a-d, Distribution of B16F10 cells and C57BL/6 mice fusion genes in chromosomes. e-h, Distribution of CT26 cells and BALB/c mice fusion genes in chromosomes.

**Table1.**
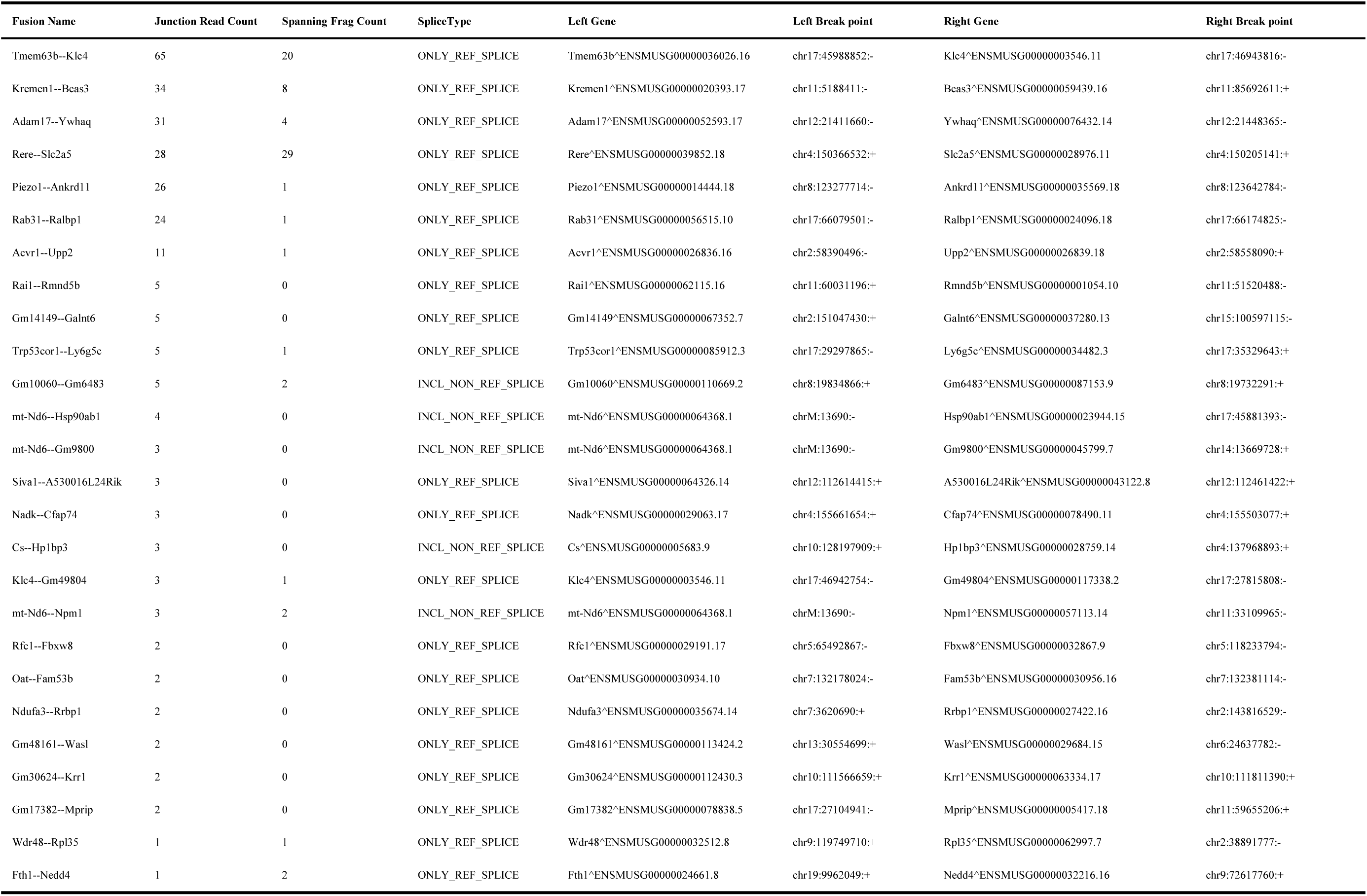
B16F10 Fusion Genes.

**Table2.**
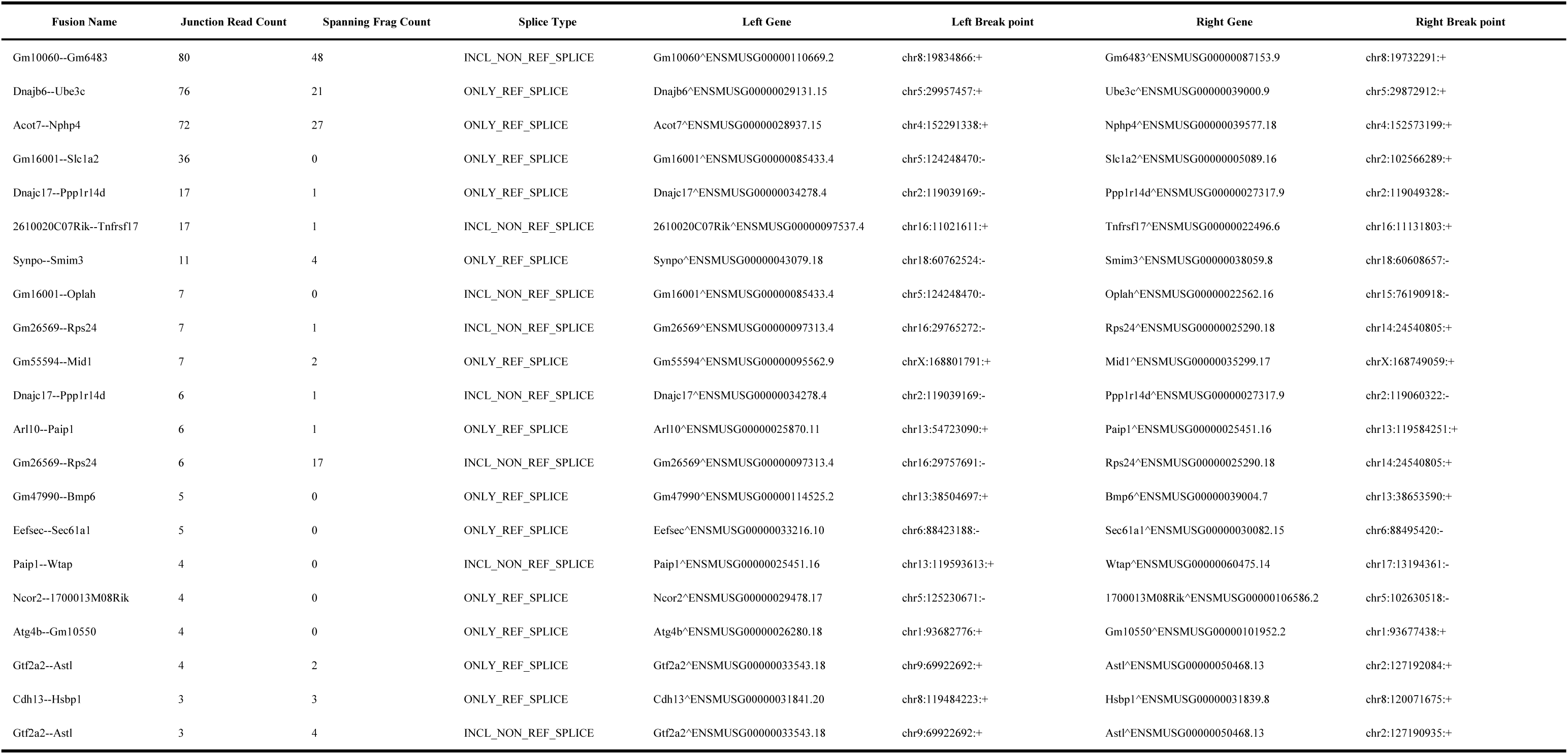
CT26 Fusion Genes.

Based on the discovery that fusion genes were extensive existed in chromosomes, we investigated the distribution of mutated locus on chromosomes. The SNP (Single Nucleotide Polymorphism) and INDELS (Insertion and Deletion) category of genome mutation were shown in brown and green respectively, and distributed in different parts of the chromosomes(figure 9 a & b). Alternative splicing is one regulatory mechanism of gene expression that a single gene can generate multiple mRNA isoforms by making different exons combinations^[30]^. Alternative splicing significantly contributes to the transcriptome complexity. By aligning RNA-seq reads to the reference genome, we identified splice junctions and analyzed the alternative splicing events of B16F10 and CT26 cells (figure S3, 9 c & d). Alternative splicing events of B16F10 and CT26 cells were mainly classified into 8 basic types, which including cassette: skipped exon, mutually exclusive exon (cassette_multi), alternative 5’ splice site (A5SS), alternative 3’ splice site (A3SS), alternative first exon (AltStart), alternative last exon (AltEnd), retained intron (IR), Mutually exclusive exons (MXE). The most common category of alternative splicing events are skipped exon. Exon skipping can result in loss of functional domains/sites or frame shifting of the open reading frame. Mgll and CD44 were found cassette exon skipping in B16F10 cells (supplementary materials), which may affect the fatty acids metabolism and adheren junction functions. MXE splicing is the only type of alternative splicing that can maintain the near same size of the encoding protein^[31]^. MBP gene was found one MXE type of alternative splicing in tumor cells (supplementary materials), which may affect the myelin sheath formation in the nervous system. This information of alternative splicing allows for the quantification and characterization of transcript isoforms, providing insights into isoform diversity, tissue-specific expression, and therapeutic target selection.

**Figure 9.**
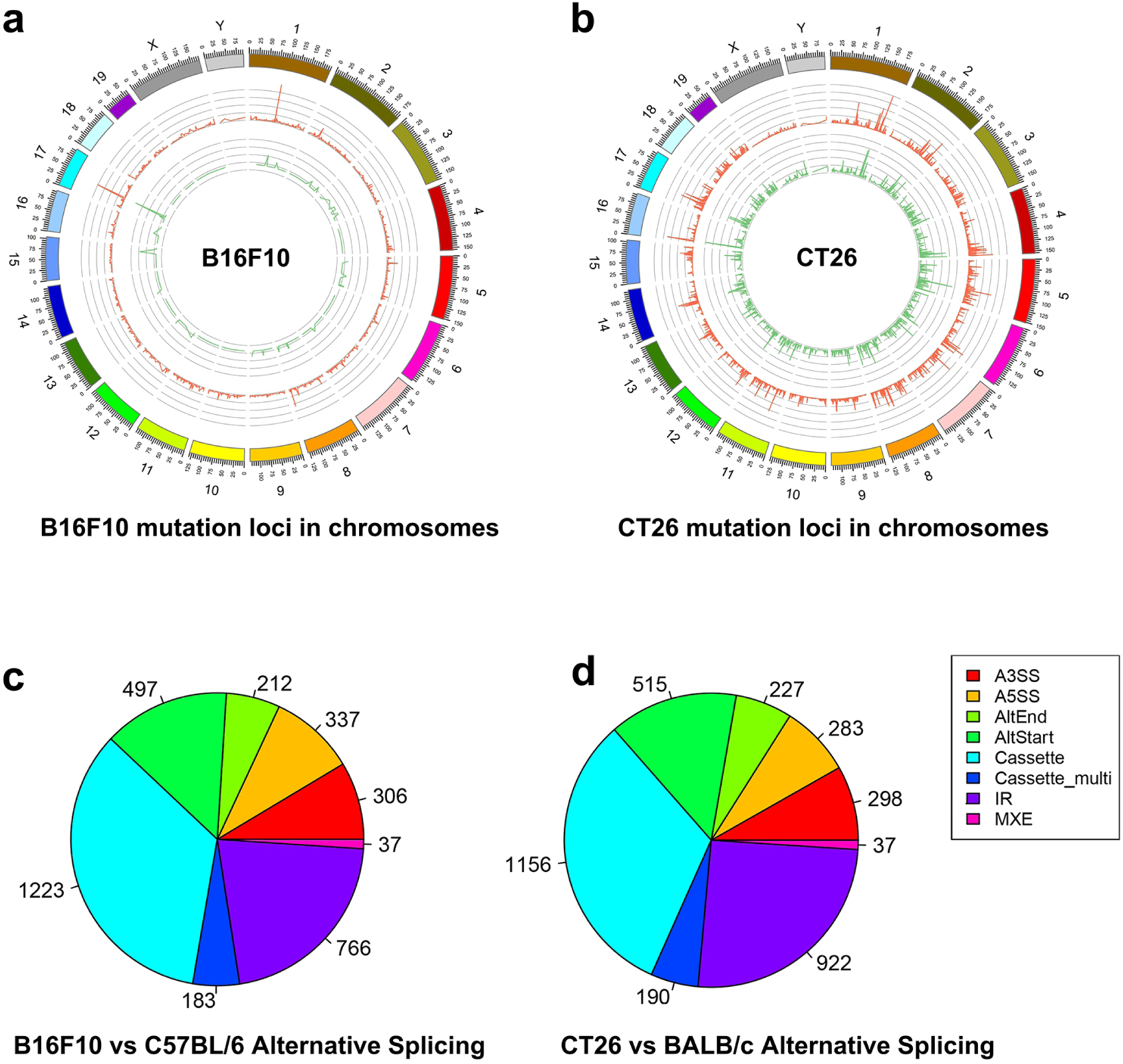
Distribution of mutation loci in chromosomes and alternative splicing. a, Distribution of B16F10 cells mutation loci on chromosome. b, Distribution of CT26 cells mutation loci on chromosome. c, The alternative splicing categories of B16F10 cells vs C57BL/6 mice. d. The alternative splicing categories of CT26 cells vs BALB/c mice. A3SS: alternative 3’ splice site, A5SS: alternative 5’ splice site, AltStart: alternative first exon, AltEnd: alternative last exon, IR: retained intron, MXE: Mutually exclusive exons

## Discussion

Next-generation sequencing (NGS) is a massively parallel sequencing technology in biological research. NGS can be used to determine the nucleotides sequence of DNA or RNA in a high-throughput and cost-effective manner, allowing comprehensive exploration of the genetic landscape of diseases^[32]^. Since mRNA coding the final expressed proteins, we utilized RNA-Seq to investigate the transcriptomics of B16F10 and CT26 tumor cells. RNA-Seq study involves the sequencing and quantification of mRNA molecules, providing a comprehensive illustration of the expressed genes. By generating millions of sequencing gene reads of tumor cells and normal tissues, RNA-Seq facilitated the identification of DEGs at the transcriptome level. The RNA-seq data was analyzed to compare differential gene expression between tumor cells and normal tissues, discover novel transcripts, assess alternative splicing events, and allow us to gain deep insights into gene expression, non-coding RNA regulation, and various biological processes and diseases.

Tumor cell lines are powerful tools for studying cancer biology, validating cancer targets and defining drug efficacy. Conventionally, cell lines can maintain the stability of genome and transcriptome, while cell lines are also sensitive to the subtle change of external environment, such as the culture media and condition^[33]^. Large panels of comprehensively characterized human cancer cell lines, including the Cancer Cell Line Encyclopedia (CCLE), have provided a rigorous backbone upon which to study genetic variants, candidate targets and therapeutic agents for cancer study^[34]^. B16F10 and CT26 cell lines are widely used for preclinical tumor mouse models establishment. The two tumor mouse models are characterized with relatively cold immune environment and resistant to immune checkpoint blockade therapy. DEGs analysis reveled that more up-regulated genes were enriched in B16F10 and CT26 tumor cells. The DEGs are mainly associated with the processing of cell cycle and division, including DNA replication, mRNA processing, organelle localization, *etc*. The results reveled the tumor cell lines have potential advantages in division and proliferation over normal tissue cells in the transcriptome level.

Fusion gene is a hallmark of many solid tumors and hematopoietic malignancies^[35]^. mRNA splicing and chromosomal rearrangements that concatenate two different genes together can form a fusion gene. Tumor cells are highly genomic instability, and fusions can occur as a result of chromosome rearrangements including translocation, insertion, inversion, deletion, tandem duplication, and chromothripsis^[36]^. Sine genes and RNA fusions tend to facilitate the tumor growth and metastasis, some fusion genes have been identified as biomarkers and prognostic factors in patients with tumors. Such as BCR-ABL1 fusion has been widely used as a biomarker and prognostic factor in patients with ALL^[37]^. EML4-ALK fusion gene is one of the most important pathogenic driver genes of NSCLC and new generation of EML4-ALK-targeted therapy is exploiting to improve patients’ prognoses^[38]^. However, although many fusion genes have been identified and reported as biomarkers of various types of tumors, fusion genes are also discovered in normal cells. For example, JAZF1-SUZ12 gene fusion exists in endometrial stromal sarcomas and normal endometrial cells^[39]^. The fusion gene may have important functions in normal physiology. In this research, we identified that fusion genes of B16F10 and CT26 cells were widely existed within one chromosome or between chromosomes.

Cancer is a disease that some mutational cells grow uncontrollably and spread to other parts of the body. The ideal precision medicine in cancer treatment is to discover the therapeutic strategies that specifically target cancer cells without affecting normal cells. Now NGS technology facilitates identification of tumor mutations with high immunogenicity, drug-resistance and perpetual proliferation potential. DEGs, fusion genes and alternative splicing analysis in this study help us understand the physiological change and mechanisms of sustaining proliferation of B16F10 and CT26 cells. By analysing the transcriptional DEGs of B16F10 and CT26 cells, we revealed that the critical DEGs of tumor cells were highly associated with the cell cycle and division. Analyzing the transcriptome patterns can shed light on the phenotypic changes of the tumor cells as well as on the dynamics of cellular behavior associated with transcriptomic changes. The analytical methods can be further utilized for targeted and personalized treatment in the future.

## Methods

### Animal

8-10-week-old female C57BL/6 mice were purchased from Shanghai Model Organisms Center. All mice were maintained in a specific pathogen-free facility at the Experimental Animal Center of Akeso Biopharma. All animal experiments were performed according to the Ministry of Health national guidelines for housing and care of laboratory animals and approved by the Akeso Biopharma Animal Experiment Ethics Committee (LAW-2024-056).

### High-throughput transcriptome sequencing (RNA-Seq)

B6F10 cells, CT26 cells and three C57BL/6 & BALB/c mouse tail tip were collected for RNA-Seq. Briefly, Total mRNA of cells and tissues was isolated using oligo(dT) magnetic beads (Thermo Scientific). Fragmented mRNA was converted to cDNA using random primers and SuperScriptll (Invitrogen) followed by second strand synthesis using DNA polymerase I and RNaseH. cDNA was end repaired using T4 DNA polymerase, Klenow DNA polymerase and 5’ phosphorylated using T4 polynucleotide kinase. Blunt ended cDNA fragments were 3’ adenylated using Klenow fragment (3’ to 5’ exo minus). 3’ single T- overhang Illumina multiplex specific adapters were ligated using a 10: 1 molar ratio of adapter to cDNA insert using T4 DNA ligase. cDNA libraries were purified and size selected at 200-220 bp using the E-Gel 2% SizeSelect gel (Invitrogen). All cleanups were done using Agencourt AMPure XP magnetic beads. Barcoded RNA-Seq libraries were clustered on the cBot using Truseq SR cluster kit v2.5 and sequenced on the Illumina HiSeq2000.

### Quality control

Raw data (raw reads) of fastq format were processed with fastp, an ultra-fast FASTQ preprocessor with useful quality. Clean data (clean reads) were obtained after quality control, adapter trimming, quality filtering and per-read quality cutting. All the downstream analyses were based on the clean data with high quality.

### Reads mapping to the reference genome

Reference genome and gene model annotation files were downloaded from genome website directly. Paired-end clean reads were aligned to the reference genome using HISAT2 v2.1.0 (hierarchical indexing for spliced alignment of transcripts), which is a highly efficient system for aligning reads from RNA sequencing experiments. HISAT2 uses an indexing scheme based on the Burrows-Wheeler transform and the Ferragina-Manzini (FM) index, employing two types of indexes for alignment: a whole genome FM index to anchor each alignment and numerous local FM indexes for very rapid extensions of these alignments. HISAT2 is the fastest system currently available, with equal or better accuracy than any other method. HISAT2 was run with the default parameters.

### Differential expression genes (DEGs) analysis

Differential expression analysis of two conditions/groups (two biological replicates per condition) was performed using the DESeq R package (1.18.1). DESeq provide statistical routines for determining differential expression in digital gene expression data using a model based on the negative binomial distribution. The resulting P-values were adjusted using the Benjamini and Hochberg’s approach for controlling the false discovery rate. Genes with |log2(FoldChange)| > 1 and adjusted P value <0.05 found by DESeq were assigned as differentially expressed.

### GO and KEGG enrichment analysis of DEGs

Gene Ontology (GO) and KEGG enrichment analysis and visualization of differentially expressed genes was implemented by the clusterProfiler R package, which is a simple-to-use tool to analyze high-throughput data obtained from transcriptomics or proteomics. Genes with adjusted Pvalue less than 0.05 were considered significantly enriched by differential expressed genes.

### Alternative splicing analysis

Alternative splicing events were classified to 7 basic types by the software CASH (Comprehensive alternative splicing hunting). CASH^[10]^ is a user-friendly software that aims to recognize and detect differential alternative splicing(AS) exons in two samples of RNA-Seq data.

### Variant calling and annotation

Bcftools v1.12 were used to perform SNP and Indel calling. ANNOVAR was used for functional annotation of variants.

### Fusion Gene Detection

Fusion genes were detected by STAR-Fusion^[40]^, a method that is both fast and accurate in identifying fusion transcripts from RNA-Seq data. STAR-Fusion can performs a fast mapping of fusion evidence to reference transcript structure annotations and filters likely artifacts to report accurate fusion predictions.

## Abbreviations

NGS: Next Generation Sequencing
DEGs: Differentially Expressed Genes
WES: Whole Exome Sequencing
WGS: Whole Genome Sequencing
GWAS: Genome Wide Association Studies
RNA-Seq: mRNA sequencing
PCA: Principal component analysis
GO: Gene Ontology.
KEGG: Kyoto Encyclopedia of Genes and Genomes
BP: Biological Process
CC: Cellular Component
MF: Molecular Function
CCLE: Cancer Cell Line Encyclopedia
TPM: Transcripts Per Kilobase of exon model per Million mapped reads
SNP: Single Nucleotide Polymorphism
INDELS: Insertion and Deletion
CASH: Comprehensive alternative splicing hunting
A5SS: 5’ splice site
A3SS: alternative 3’ splice site
AltStart: alternative first exon
AltEnd: alternative last exon
IR: retained intron
MXE: Mutually exclusive exons

## Ethics statement

The animal study was reviewed and approved by Akeso Biopharma Animal Experiment Ethics Committee(LAW-2024-056).

## Author contributions

XZ and YF designed the study. XZ drafted the manuscript. All authors were involved in manuscript preparation and revisions. All authors read and approved the final manuscript.

## Availability of data and materials

All data and materials are included in the manuscript and supplmently materials.

## Reference

[1] Stevens JB, Horne SD, Abdallah BY, Ye CJ, Heng HH. Chromosomal instability and transcriptome dynamics in cancer. Cancer Metastasis Rev. 2013 Dec;32(3-4):391–402. doi: 10.1007/s10555-013-9428-6. PMID: 23595307.

[2] Pon JR, Marra MA. Driver and passenger mutations in cancer. Annu Rev Pathol. 2015;10:25–50. doi: 10.1146/annurev-pathol-012414-040312. Epub 2014 Oct 17. PMID: 25340638.

[3] Hong M, Tao S, Zhang L, Diao LT, Huang X, Huang S, Xie SJ, Xiao ZD, Zhang H. RNA sequencing: new technologies and applications in cancer research. J Hematol Oncol. 2020 Dec 4;13(1):166. doi: 10.1186/s13045-020-01005-x. PMID: 33276803; PMCID: PMC7716291.

[4] Hakimi AA, Voss MH, Kuo F, Sanchez A, Liu M, Nixon BG, Vuong L, Ostrovnaya I, Chen YB, Reuter V, Riaz N, Cheng Y, Patel P, Marker M, Reising A, Li MO, Chan TA, Motzer RJ. Transcriptomic Profiling of the Tumor Microenvironment Reveals Distinct Subgroups of Clear Cell Renal Cell Cancer: Data from a Randomized Phase III Trial. Cancer Discov. 2019 Apr;9(4):510–525. doi: 10.1158/2159-8290.CD-18-0957. Epub 2019 Jan 8. PMID: 30622105; PMCID: PMC6697163.

[5] Larson NB, Oberg AL, Adjei AA, Wang L. A Clinician’s Guide to Bioinformatics for Next-Generation Sequencing. J Thorac Oncol. 2023 Feb;18(2):143–157. doi: 10.1016/j.jtho.2022.11.006. Epub 2022 Nov 12. PMID: 36379355; PMCID: PMC9870988.

[6] Rusch M, Nakitandwe J, Shurtleff S, Newman S, Zhang Z, Edmonson MN, Parker M, Jiao Y, Ma X, Liu Y, Gu J, Walsh MF, Becksfort J, Thrasher A, Li Y, McMurry J, Hedlund E, Patel A, Easton J, Yergeau D, Vadodaria B, Tatevossian RG, Raimondi S, Hedges D, Chen X, Hagiwara K, McGee R, Robinson GW, Klco JM, Gruber TA, Ellison DW, Downing JR, Zhang J. Clinical cancer genomic profiling by three-platform sequencing of whole genome, whole exome and transcriptome. Nat Commun. 2018 Sep 27;9(1):3962. doi: 10.1038/s41467-018-06485-7. PMID: 30262806; PMCID: PMC6160438.

[7] Satam H, Joshi K, Mangrolia U, Waghoo S, Zaidi G, Rawool S, Thakare RP, Banday S, Mishra AK, Das G, Malonia SK. Next-Generation Sequencing Technology: Current Trends and Advancements. Biology (Basel). 2023 Jul 13;12(7):997. doi: 10.3390/biology12070997. Erratum in: Biology (Basel). 2024 Apr 24;13(5):286. doi: 10.3390/biology13050286. PMID: 37508427; PMCID: PMC10376292.

[8] Wu Y, Zheng Z, Visscher PM, Yang J. Quantifying the mapping precision of genome-wide association studies using whole-genome sequencing data. Genome Biol. 2017 May 16;18(1):86. doi: 10.1186/s13059-017-1216-0. PMID: 28506277; PMCID: PMC5432979.

[9] Konda P, Garinet S, Van Allen EM, Viswanathan SR. Genome-guided discovery of cancer therapeutic targets. Cell Rep. 2023 Aug 29;42(8):112978. doi: 10.1016/j.celrep.2023.112978. Epub 2023 Aug 10. PMID: 37572322.

[10] Chen S, Jiang W, Du Y, Yang M, Pan Y, Li H, Cui M. Single-cell analysis technologies for cancer research: from tumor-specific single cell discovery to cancer therapy. Front Genet. 2023 Oct 12;14:1276959. doi: 10.3389/fgene.2023.1276959. PMID: 37900181; PMCID: PMC10602688.

[11] Du P, Fan R, Zhang N, Wu C, Zhang Y. Advances in Integrated Multi-omics Analysis for Drug-Target Identification. Biomolecules. 2024 Jun 14;14(6):692. doi: 10.3390/biom14060692. PMID: 38927095; PMCID: PMC11201992.

[12] Moyerbrailean GA, Davis GO, Harvey CT, Watza D, Wen X, Pique-Regi R, Luca F. A high-throughput RNA-seq approach to profile transcriptional responses. Sci Rep. 2015 Oct 29;5:14976. doi: 10.1038/srep14976. PMID: 26510397; PMCID: PMC4625130.

[13] Han Y, Gao S, Muegge K, Zhang W, Zhou B. Advanced Applications of RNA Sequencing and Challenges. Bioinform Biol Insights. 2015 Nov 15;9(Suppl 1):29–46. doi: 10.4137/BBI.S28991. PMID: 26609224; PMCID: PMC4648566.

[14] Byron SA, Van Keuren-Jensen KR, Engelthaler DM, Carpten JD, Craig DW. Translating RNA sequencing into clinical diagnostics: opportunities and challenges. Nat Rev Genet. 2016 May;17(5):257–71. doi: 10.1038/nrg.2016.10. Epub 2016 Mar 21. PMID: 26996076; PMCID: PMC7097555.

[15] Wang Z, Gerstein M, Snyder M. RNA-Seq: a revolutionary tool for transcriptomics. Nat Rev Genet. 2009 Jan;10(1):57–63. doi: 10.1038/nrg2484. PMID: 19015660; PMCID: PMC2949280.

[16] Ozsolak F, Milos PM. RNA sequencing: advances, challenges and opportunities. Nat Rev Genet. 2011 Feb;12(2):87–98. doi: 10.1038/nrg2934. PMID: 21191423; PMCID: PMC3031867.

[17] Liang P, Pardee AB. Analysing differential gene expression in cancer. Nat Rev Cancer. 2003 Nov;3(11):869–76. doi: 10.1038/nrc1214. PMID: 14668817.

[18] Ben-David U, Siranosian B, Ha G, Tang H, Oren Y, Hinohara K, Strathdee CA, Dempster J, Lyons NJ, Burns R, Nag A, Kugener G, Cimini B, Tsvetkov P, Maruvka YE, O’Rourke R, Garrity A, Tubelli AA, Bandopadhayay P, Tsherniak A, Vazquez F, Wong B, Birger C, Ghandi M, Thorner AR, Bittker JA, Meyerson M, Getz G, Beroukhim R, Golub TR. Genetic and transcriptional evolution alters cancer cell line drug response. Nature. 2018 Aug;560(7718):325–330. doi: 10.1038/s41586-018-0409-3. Epub 2018 Aug 8. PMID: 30089904; PMCID: PMC6522222.

[19] Ben-David U, Siranosian B, Ha G, Tang H, Oren Y, Hinohara K, Strathdee CA, Dempster J, Lyons NJ, Burns R, Nag A, Kugener G, Cimini B, Tsvetkov P, Maruvka YE, O’Rourke R, Garrity A, Tubelli AA, Bandopadhayay P, Tsherniak A, Vazquez F, Wong B, Birger C, Ghandi M, Thorner AR, Bittker JA, Meyerson M, Getz G, Beroukhim R, Golub TR. Genetic and transcriptional evolution alters cancer cell line drug response. Nature. 2018 Aug;560(7718):325–330. doi: 10.1038/s41586-018-0409-3. Epub 2018 Aug 8. PMID: 30089904; PMCID: PMC6522222.

[20] Gillet JP, Varma S, Gottesman MM. The clinical relevance of cancer cell lines. J Natl Cancer Inst. 2013 Apr 3;105(7):452–8. doi: 10.1093/jnci/djt007. Epub 2013 Feb 21. PMID: 23434901; PMCID: PMC3691946.

[21] Huang L, Li Y, Du Y, Zhang Y, Wang X, Ding Y, Yang X, Meng F, Tu J, Luo L, Sun C. Mild photothermal therapy potentiates anti-PD-L1 treatment for immunologically cold tumors via an all-in-one and all-in-control strategy. Nat Commun. 2019 Oct 25;10(1):4871. doi: 10.1038/s41467-019-12771-9. PMID: 31653838; PMCID: PMC6814770.

[22] Sato Y, Fu Y, Liu H, Lee MY, Shaw MH. Tumor-immune profiling of CT-26 and Colon 26 syngeneic mouse models reveals mechanism of anti-PD-1 response. BMC Cancer. 2021 Nov 13;21(1):1222. doi: 10.1186/s12885-021-08974-3. PMID: 34774008; PMCID: PMC8590766.

[23] Sims D, Sudbery I, Ilott NE, Heger A, Ponting CP. Sequencing depth and coverage: key considerations in genomic analyses. Nature reviews. Genetics. 2014 Feb;15(2):121–132. DOI: 10.1038/nrg3642. PMID: 24434847.

[24] Croft D, O’Kelly G, Wu G, Haw R, Gillespie M, Matthews L, Caudy M, Garapati P, Gopinath G, Jassal B, Jupe S, Kalatskaya I, Mahajan S, May B, Ndegwa N, Schmidt E, Shamovsky V, Yung C, Birney E, Hermjakob H, D’Eustachio P, Stein L. Reactome: a database of reactions, pathways and biological processes. Nucleic Acids Res. 2011 Jan;39(Database issue):D691–7. doi: 10.1093/nar/gkq1018. Epub 2010 Nov 9. PMID: 21067998; PMCID: PMC3013646.

[25] Yu G, Wang LG, Han Y, He QY. clusterProfiler: an R package for comparing biological themes among gene clusters. OMICS. 2012 May;16(5):284–7. doi: 10.1089/omi.2011.0118. Epub 2012 Mar 28. PMID: 22455463; PMCID: PMC3339379.

[26] Wang L, Yang H, Hu L, Hu D, Ma S, Sun X, Jiang L, Song J, Ji L, Masau JF, Zhang H, Qian K. CDKN1C (P57): one of the determinants of human endometrial stromal cell decidualization. Biol Reprod. 2018 Mar 1;98(3):277–285. doi: 10.1093/biolre/iox187. PMID: 29325014.

[27] Wang Y, Wu H, Dong N, Su X, Duan M, Wei Y, Wei J, Liu G, Peng Q, Zhao Y. Sulforaphane induces S-phase arrest and apoptosis via p53-dependent manner in gastric cancer cells. Sci Rep. 2021 Jan 28;11(1):2504. doi: 10.1038/s41598-021-81815-2. PMID: 33510228; PMCID: PMC7843980.

[28] Annala MJ, Parker BC, Zhang W, Nykter M. Fusion genes and their discovery using high throughput sequencing. Cancer Lett. 2013 Nov 1;340(2):192–200. doi: 10.1016/j.canlet.2013.01.011. Epub 2013 Jan 29. PMID: 23376639; PMCID: PMC3675181.

[29] Taniue K, Akimitsu N. Fusion Genes and RNAs in Cancer Development. Noncoding RNA. 2021 Feb 4;7(1):10. doi: 10.3390/ncrna7010010. PMID: 33557176; PMCID: PMC7931065.

[30] Chen M, Manley JL. Mechanisms of alternative splicing regulation: insights from molecular and genomics approaches. Nat Rev Mol Cell Biol. 2009 Nov;10(11):741–54. doi: 10.1038/nrm2777. Epub 2009 Sep 23. PMID: 19773805; PMCID: PMC2958924.

[31] Pohl M, Bortfeldt RH, Grützmann K, Schuster S. Alternative splicing of mutually exclusive exons--a review. Bio Systems. 2013 Oct;114(1):31–38. DOI: 10.1016/j.biosystems.2013.07.003. PMID: 23850531.

[32] Satam H, Joshi K, Mangrolia U, Waghoo S, Zaidi G, Rawool S, Thakare RP, Banday S, Mishra AK, Das G, Malonia SK. Next-Generation Sequencing Technology: Current Trends and Advancements. Biology (Basel). 2023 Jul 13;12(7):997. doi: 10.3390/biology12070997. Erratum in: Biology (Basel). 2024 Apr 24;13(5):286. doi: 10.3390/biology13050286. PMID: 37508427; PMCID: PMC10376292.

[33] Geraghty RJ, Capes-Davis A, Davis JM, Downward J, Freshney RI, Knezevic I, Lovell-Badge R, Masters JR, Meredith J, Stacey GN, Thraves P, Vias M; Cancer Research UK. Guidelines for the use of cell lines in biomedical research. Br J Cancer. 2014 Sep 9;111(6):1021–46. doi: 10.1038/bjc.2014.166. Epub 2014 Aug 12. PMID: 25117809; PMCID: PMC4453835.

[34] Ghandi M, Huang FW, Jané-Valbuena J, Kryukov GV, et al. Next-generation characterization of the Cancer Cell Line Encyclopedia. Nature. 2019 May;569(7757):503–508. doi: 10.1038/s41586-019-1186-3. Epub 2019 May 8. PMID: 31068700; PMCID: PMC6697103.

[35] Taniue K, Akimitsu N. Fusion Genes and RNAs in Cancer Development. Noncoding RNA. 2021 Feb 4;7(1):10. doi: 10.3390/ncrna7010010. PMID: 33557176; PMCID: PMC7931065.

[36] Taniue K, Akimitsu N. Fusion Genes and RNAs in Cancer Development. Noncoding RNA. 2021 Feb 4;7(1):10. doi: 10.3390/ncrna7010010. PMID: 33557176; PMCID: PMC7931065.

[37] Jain S, Abraham A. BCR-ABL1-like B-Acute Lymphoblastic Leukemia/Lymphoma: A Comprehensive Review. Arch Pathol Lab Med. 2020 Feb;144(2):150–155. doi: 10.5858/arpa.2019-0194-RA. Epub 2019 Oct 23. PMID: 31644323.

[38] Lei Y, Lei Y, Shi X, Wang J. EML4-ALK fusion gene in non-small cell lung cancer. Oncol Lett. 2022 Jun 24;24(2):277. doi: 10.3892/ol.2022.13397. PMID: 35928804; PMCID: PMC9344266.

[39] Li H, Wang J, Ma X, Sklar J. Gene fusions and RNA trans-splicing in normal and neoplastic human cells. Cell Cycle. 2009 Jan 15;8(2):218–22. doi: 10.4161/cc.8.2.7358. Epub 2009 Jan 6. PMID: 19158498.

[40] Haas BJ, Dobin A, Stransky N, et al. STAR-Fusion: Fast and Accurate Fusion Transcript Detection from RNA-Seq. bioRxiv; 2017. DOI: 10.1101/120295.

